# Mechanistic insights into volatile anesthetic modulation of K2P channels

**DOI:** 10.1101/2020.06.10.144287

**Authors:** Aboubacar Wague, Thomas T. Joseph, Kellie A. Woll, Weiming Bu, Kiran A. Vaidya, Natarajan V. Bhanu, Benjamin A. Garcia, Crina M. Nimigean, Roderic G. Eckenhoff, Paul M. Riegelhaupt

## Abstract

K2P potassium channels are known to be modulated by volatile anesthetic (VA) drugs and play important roles in clinically relevant effects that accompany general anesthesia. Here, we utilize a photoaffinity analog of the VA isoflurane to identify a VA binding site in the TREK1 K2P channel. The functional importance of the identified site was validated by mutagenesis and biochemical modification. Molecular dynamic simulations of TREK1 in the presence of VA found multiple neighboring residues on TREK1 TM2, TM3 and TM4 that contribute to anesthetic binding. The identified VA binding region contains residues that play roles in the mechanisms by which heat, mechanical stretch, and pharmacological modulators alter TREK1 channel activity and overlaps with positions found to modulate TASK K2P channel VA sensitivity. Our findings define molecular contacts that mediate VA binding to TREK1 channels and suggest a mechanistic basis to explain how K2P channels are modulated by VAs.

## Introduction

Tandem pore (K2P) potassium channels are a group of physiologically important K^+^ leak channels that modulate cellular resting membrane potential to control excitability (Enyedi and Czirjak 2010). Human K2P channel abnormalities cause a wide range of physiological syndromes, including cardiac conduction abnormalities that lead to ventricular tachycardia and fibrillation (Gurney and Manoury 2009, Friedrich et al. 2014, Decher et al. 2017), familial pulmonary arterial hypertension (Ma et al. 2013), salt-sensitive hyperaldosteronism (Davies et al. 2008), and familial migraines (Lafreniere et al. 2010, Royal et al. 2019), underscoring roles for K2P channels in human cardiac, vascular, renal, and pain physiology. K2Ps are proven contributors to the potency of volatile anesthetics (VAs), with K2P knockout animals exhibiting a measurable resistance to VA induced loss of consciousness (Heurteaux et al. 2004, Pang et al. 2009, Steinberg et al. 2015). Systemic effects of VA administration have been directly linked to K2P channel activity, including a role for carotid body TASK channels in mediating VA induced respiratory depression (Cotten 2013, Chokshi et al. 2015) and for TREK1 channels in the vasodilatory and neuroprotective effects of both polyunsaturated fatty acids and VAs (Blondeau et al. 2007, Tong et al. 2014).

Initial evidence for a VA induced potassium leak current was found in molluscan pacemaker neurons, an activity subsequently determined to be mediated by a member of the TASK K2P subfamily (Franks and Lieb 1988, Andres-Enguix et al. 2007). Follow-up studies of both snail and human TASK channels identified specific amino acids that alter TASK responsiveness to VAs or eliminate stereoselective discrimination between enantiomeric VA agents (Andres-Enguix et al. 2007, Conway and Cotten 2012, Luethy et al. 2017). These findings enabled predictions of structural determinants of anesthetic binding within the TASK channel family (Bertaccini et al. 2014). However, significant differences in the gating and regulation of divergent K2P subfamilies make it difficult to generalize molecular details of the interactions between VAs and TASK K2P channels to other anesthetic sensitive K2Ps (Patel et al. 1999), most notably the TREK1 channel.

TREK1 is a member of a subgroup of K2Ps (including TREK1, TREK2, and TRAAK) regulated by the biophysical properties of their surroundings, including membrane stretch or deformation, temperature, and bilayer lipid composition (Honore 2007). This pronounced sensitivity of TREK1 to its surrounding environment is unique amongst known biologically relevant VA targets. VAs are small hydrophobic drugs that readily partition into lipid bilayers and *in-vivo* anesthetic potency has long been known to correlate with lipid solubility, a relationship that holds true over a wide range of structurally diverse anesthetic agents (Sonner and Cantor 2013). While the majority of studies exploring the interaction between anesthetics and ion channels have identified discrete binding sites thought to underlie the modulatory effects of these drugs (Nury et al. 2011, Sauguet et al. 2013, Hemmings et al. 2019), modulation of TREK1 by VAs could be posited to occur via indirect effects on the properties of the surrounding bilayer. In fact, a recent study has proposed that in vitro administration of VA agents disrupt membrane lipid rafts, altering phospholipase C activity and modifying the milieu of lipids surrounding TREK1 to modulate its activity (Pavel et al. 2019).

The question of how VA drugs modulate TREK1 is the key issue addressed in this study. By utilizing an unbiased photolabeling approach, we show that isoflurane, a volatile anesthetic, binds to a TREK1 TM2 residue in close proximity to the previously proposed VA modulatory site in TASK K2P channels. MD simulation studies based on our photolabeling findings show that isoflurane, a known low affinity ligand for TREK1, is mobile within this newly identified TREK1 binding site and interacts with residues from TM2, TM3, and TM4. These interactions include key molecular contacts with resides known to influence the conformational rearrangement of the TM4 helix that drive K2P modulation by heat, mechanical stretch, and other pharmacological channel modulators (Dong et al. 2015, Lolicato et al. 2014, Brohawn, Campbell and MacKinnon 2014a, Pope et al. 2018). Our photolabeling and MD results enabled us to design multiple point mutations within the putative VA binding pocket that specifically diminished TREK1 VA responsiveness without otherwise perturbing channel function. Our data support the notion of a shared VA modulatory site across the TREK and TASK K2P subfamilies and suggest a mechanism by which binding of VA within this site promotes conformational rearrangements known to gate K2P channels.

## Experimental Methods

### Purification of drTREK1 or hTRAAK

TREK1 and TRAAK proteins were expressed in *Pichia pastoris* using previously described pPICZ vectors bearing residues 1–300 of the human TRAAK gene (K2P4.1, UniProt Q9NYG8-2) or residues 1–322 of the *D. rerio* TREK1 gene (K2P2.1, UniProt X1WC65), followed by a PreScission protease-cleavage site (LEVLFQ/GP) and C-terminal GFP and His10 tags (Brohawn, Su and MacKinnon 2014b). Mutations to eliminate N-linked glycosylation sites were inserted into both the human TRAAK (N104Q and N108Q) and zebrafish TREK1 (N95Q, N122Q) gene constructs. Expression plasmids were linearized with the PmeI restriction enzyme and were subsequently transformed by electroporation into *P. pastoris* strain SMD1168H. Screening for successful recombinant integration was performed by plating transformants on yeast extract peptone dextrose sorbitol (YPDS) plates containing increasing concentrations of zeocin from 0.5 to 3 mg/ml.

Screening, expression, and purification of K2Ps was performed as previously described (Lolicato et al. 2014). Briefly, yeast transformants were grown in buffered minimal medium (2 × YNB, 1% glycerol, 0.4 mg L^−1^ biotin, 100 mM potassium phosphate [pH 6.0]) for 2 days at 30°C in a shaker at 225 rpm. Cells were then pelleted by centrifugation (4,000 × g at 20°C, 5 min) and resuspended in methanol minimal medium (2 × YNB, 0.5% methanol, 0.4 mg L^−1^ biotin, 100 mM potassium phosphate [pH 6.0]) to induce protein expression. Cells were then shaken for 2 additional days at 22°C in a shaker at 225 rpm, with additional methanol added (final concentration 0.5% [v/v]) to the culture after 24 hours of protein expression. After 48 hours of protein expression, cells were pelleted by centrifugation (6,000 × g at 4°C for 10 min), flash frozen in liquid nitrogen, and then subjected to three rounds of cryo-milling (Retsch model MM301) in liquid N_2_ for 3 min at 25 Hz to disrupt yeast cell walls and membranes. Frozen yeast cell powder was then stored at −80°C.

Cell powder was added to breaking buffer (150 mM KCl, 50 mM Tris pH 8.0, 1 mM phenylmethysulfonyl fluoride, 0.1 mg/mL DNase 1, and 1 tablet/50 ml of EDTA-free complete inhibitor cocktail [Roche]) at a ratio of 1 g cell pellet/4 mL lysis buffer. Solubilized cell powder was centrifuged at 4,000 × g at 4°C for 5 min to pellet large debris and the supernatant was then centrifuged at 100,000 × g at 4°C for 1.5 hours to pellet cell membranes. The pellet was re-suspended in 50 ml breaking buffer containing 60mM n-Dodecyl-B-D-Maltoside (DDM) and incubated for 3 hours with gentle stirring to solubilize the membranes, followed by centrifugation at 35,000 × g for 45 min. Talon cobalt resin (Takara Bio USA) was added to the supernatant at a ratio of 1 ml of resin per 10 g of cell powder and incubated in an orbital rotor overnight at 4°C. Resin was then collected on a column and washed with 10 column volumes of Buffer A (150 mM KCl, 50 mM Tris pH 8.0, 6 mM DDM, 30 mM imidazole) and bound protein was subsequently eluted from the resin by washing with Buffer B (150 mM KCl, 50 mM Tris pH 8.0, 6 mM DDM, 300 mM imidazole). PreScission protease (~1:25 wt:wt) was added to the eluate and the cleavage reaction was allowed to proceed overnight at 4°C under gentle rocking. Cleaved TREK1 or TRAAK protein was concentrated in 50 kDa molecular weight cutoff (MWCO) Amicon Ultra Centrifugal Filters (Millipore) and applied to a Superdex200 10/300 gel filtration column (GE Healthcare) equilibrated in size exclusion chromatography (SEC) buffer (150 mM KCl, 20 mM Tris pH 8.0, 1mM DDM). Purified *h*TRAAK or *dr*TREK1 protein was concentrated (50 kDa MWCO) to 8 mg/mL prior to reconstitution and analyzed for purity by SDS-PAGE [12% (wt/vol) gels; Bio-Rad] followed by staining with coomassie blue. All protein purification steps were carried out at 4 °C.

### Reconstitution of K2P channels into liposomes

Immediately following purification, *dr*TREK1 or *h*TRAAK channel protein was reconstituted into liposomes (Heginbotham, Kolmakova-Partensky and Miller 1998). 5mg of a 3:1 (wt/wt) ratio of 1-palmitoyl-2-oleoyl-sn-glycero-3-phosphoethanolamine (POPE) and 1-palmitoyl-2-oleoyl-sn-glycero-3-phospho- (1′-rac-glycerol) (POPG) in chloroform was used for reconstitution. The lipids were dried in a borosilicate glass vial under nitrogen flow and then solubilized in reconstitution buffer (400 mM KCl, 10 mM HEPES, 5 mM NMDG [N-methylglucamine D-gluconate], pH 7.6) containing 34 mM CHAPS (3-((3-cholamidopropyl) dimethylammonio)-1-propanesulfonate). The lipid containing solution was bath sonicated until clear. *dr*TREK1 or *h*TRAAK protein was then added to the lipid solution at a concentration of 100 μg protein per mg of solubilized lipids. Proteoliposomes were formed by applying the lipid/protein containing sample (500 μl total volume) to an 18 ml detergent-removal column (Sephadex G-50 fine beads, GE Healthcare Life Sciences). Turbid fractions containing proteoliposomes were pooled, aliquoted, flash frozen in liquid nitrogen, and stored at −80°C.

### Photoaffinity labeling of drTREK1 or hTRAAK potassium channels for protein microsequencing

5-7 μg of *dr*TREK or *h*TRAAK in proteoliposomes was added to a 30 μM solution of azi-isoflurane ± 3 mM isoflurane. Each sample was equilibrated on ice in the dark for 5 min and then transferred to a 1 mm path length quartz cuvette and exposed for 25 min to 300 nm ultraviolet light produced by an RPR-3000 Rayonet lamp filtered by a WG295 295nm glass filter (Newport Corporation).

### In-Solution Protein Digestion

Photolabeled samples underwent dialysis and buffer exchange using 10 kDa MWCO Amicon Ultra Centrifugal Filters (Millipore). ProteaseMAX™ Surfactant (Promega) was added to a concentration of 0.2% and the samples were vortexed vigorously for 30 sec. Samples were then diluted with NH_4_HCO_3_ to a final concentration of 50 mM NH_4_HCO_3_ in a 93.5 μL volume. 1 μL of 0.5 M dithiothreitol (DTT) was then added and samples were incubated at 56 °C for 30 min. 2.7 μL of 0.55 M iodoacetamide (IAA) was subsequently added and protein samples were incubated at room temperature in the dark for 45 min. An additional 1 μL of 1% (w/v%) ProteaseMax ™ Surfactant was added to the sample, followed by sequencing grade-modified trypsin (Promega) to a 1:20 protease:protein final ratio (w:w). Proteins were digested overnight at 37°C. Trypsin digested peptides were again diluted with NH_4_HCO_3_ to a final concentration of 100 mM NH_4_HCO_3_ and 0.02% ProteaseMAX Surfactant in a 200 μL total volume, prior to addition of sequencing grade chymotrypsin (Promega) to a final 1:20 protease:protein ratio (w:w). Proteins were again digested overnight at 37°C. Acetic acid was added to reach a pH < 2 and the peptide digests were then incubated at room temperature for 10 min, prior to centrifugation at 16, 000 × g for 20 min to remove insoluble debris. Samples were then desalted using C18 stage tips prepared in house, dried by speed-vac, and resuspended in 0.1% formic acid immediately prior to mass spectrometry analysis.

### In-Gel Protein Digestion

Photolabeled samples underwent dialysis and buffer exchange using 10 kDa MWCO Amicon Ultra Centrifugal Filters (Millipore). Samples were then mixed with SDS loading buffer containing DTT to a final concentration of 100 mM DTT, vortexed vigorously and incubated at room temperature for 45 min before separation by SDS-PAGE. The resulting gels were stained with Coomassie Blue G250 (BioRad), destained, and washed with ddH_2_O. Protein bands between ~30-40kDa (corresponding to *dr*TREK or *h*TRAAK) were identified and excised. Excised bands were destained, dehydrated and dried by speed vac. Proteins were then reduced by incubation at 56° C for 30 min in 5 mM DTT and 50 mM NH_4_HCO_3_. The DTT solution was removed and proteins were then alkylated by the addition of 55 mM IAA in 50 mM NH_4_HCO_3_ and incubation at room temperature for 45 min in the dark. Bands were dehydrated and dried by speed vac before resuspension in 100 μL 0.2 % ProteaseMAX™ surfactant and 50 mM NH_4_HCO_3_ solution containing trypsin at a 1:20 protease:protein ratio (w:w). Proteins were digested overnight at 37°C. To enhance sequence coverage, a second protease digestion was performed with chymotrypsin. Samples were diluted to a final volume of 200 μL with final concentrations of 100 mM NH_4_HCO_3_ and approximately 0.02% ProteaseMAX™ Surfactant. Sequencing grade chymotrypsin (Promega) to a 1:20 protease:protein ratio (w:w) was then added and proteins were digested overnight at 37 °C. Multiple extraction steps were performed to remove peptides embedded in the gel. The digest solution was removed and the remaining gel was suspended in 100 μL 30% acetylnitrile and 5% acetic acid in ddH_2_O (v/v%) and sonicated for 20 min. The solution was removed and the gel was then resuspended in 100 μL 70% acetylnitrile and 5% acetic acid in ddH_2_O (v/v%) and sonicated for 20min. The digest and two extraction solutions were pooled and dried by speed vac before resuspension in 0.5% acetic acid (pH < 2). Samples were sonicated for 10 min prior to centrifugation to remove insoluble debris. Samples were desalted using C18 stage tips prepared in house. Samples were dried by speed vac and resuspended in 0.1% formic acid immediately before mass spectrometry analysis.

### Mass spectrometry

Desalted peptides were analyzed employing either an Orbitrap Elite™ Hybrid Ion Trap-Orbitrap mass spectrometer (MS) or Q Exactive™ Hybrid Quadrupole-Orbitrap MS coupled to an Easy-nanoLC 1000 system. In both instances the same liquid chromatography procedure and data dependent acquisition mode was applied. Peptides were eluted over 100 min with linear gradients of ACN in 0.1% formic acid in water (v/v%) starting from 2% to 40% (85 min), then 40% to 85% (5 min) and finally 85% (10 min) using a flow rate of 300mL/min. For Orbitrap Elite™, in every 3 s cycle, one full MS scan was collected at a scan range of 350 to 1500 m/z, a resolution of 60K, and a maximum injection time of 50 ms with automatic gain (AG) control of 500000. The MS2 scans were followed from the most intense parent ions. Ions were filtered with charge 2-5 with an isolation window of 1.5 m/z in quadruple isolation mode. Ions were fragmented using collision induced dissociation (CID) with collision energy of 35%. Iontrap detection was used with normal scan range mode and rapid iontrap scan rate. For Q Exactive™, one full MS was performed with 70K resolution, maximum injection time of 100 ms and scan range of 350-1200 m/z. The MS2 scans were performed in Higher-energy Collisional Dissociation (HCD) with normalized collision energy (NCE) of 30, and isolation window of 3 m/z. The AG control was set to be 10000 with a maximal injection time of 100 ms.

### Mass spectrometry analysis

Analysis was performed as previously reported (Eckenhoff et al., 2010; Woll et al., 2017). Spectral analysis was conducted using Thermo Proteome Discoverer 2.0 (Thermo Scientific) and the Mascot Daemon search engine with a customized database containing *dr*TREK or *h*TRAAK protein sequences. All analyses included dynamic oxidation of methionine (+15.9949 *m/z*) as well as static alkylation of cysteine (+57.0215 *m/z*; iodoacetamide alkylation). Photolabeled samples were run with the additional dynamic azi-isoflurane (+195.97143 *m/z*) modification. A mass variation tolerance of 10 ppm and 20 ppm for MS and 0.8 and 0.02 Da for MS/MS were used for the for Orbitrap Elite™ and Q Exactive™ respectively. Both the in-solution and in-gel sequential trypsin/chymotrypsin digests were searched without enzyme specification with a false discovery rate of 0.01. All MS experiments were conducted in triplicate and samples containing no photoaffinity ligand were processed and analyzed equivalently to those containing the photolabel, to control for false positive detection of photoaffinity ligand modifications.

### Molecular Dynamics Simulation

The crystallographically derived structural model of *Mus musculus* TREK1 (PDBID 6CQ6) was used for MD simulation. Missing nonterminal loops were predicted using MODELLER (Webb and Sali 2017) with side chain conformations predicted using SCRWL4 (Krivov, Shapovalov and Dunbrack 2009). Molecular mechanics parameters for isoflurane were used, as previously published (Henin et al. 2010). In the ligand-bound simulation, one isoflurane molecule was placed manually, adjacent to G182. The receptor was embedded in a POPC (1-palmitoyl-2-oleoyl-*sn*- glycero-3-phosphocholine):cholesterol 70:30 lipid bilayer and the system was solvated with TIP3P water with 0.15 M NaCl using the CHARMM-GUI service (Jo et al. 2008, Jo et al. 2009). The CHARMM36 force field (Klauda et al. 2010, Best et al. 2012) and NAMD 2.12 simulation software (Phillips et al. 2005) were used. Periodic boundary conditions with particle mesh Ewald summation of long-range electrostatics was used. The system was equilibrated according to the CHARMM-GUI protocol and production equilibrium MD simulation was run for 10 nanoseconds using the isothermic-isobaric ensemble with Langevin thermostat and barostat at 303.15 K. Further equilibrium MD simulations were continued from that point.

### Two electrode voltage clamp electrophysiology

For electrophysiological studies, *Xenopus laevis* oocytes were microinjected with capped RNA translated from full length mouse TREK1 (K2P2.1, UniProt P97438-2) or mouse TRAAK (K2P4.1, UniProt O88454) genes. TREK1 mutations were introduced using a Phusion site directed mutagenesis kit (ThermoFisher), with the TREK1 C93S C159S C219S C365S C399S mutant channel referred to as TREK1 cys-for brevity. cRNA from WT or mutant K2P genes was synthesized using an mMessage mMachine Kit (T7 promoter, Ambion, Life Technologies) and purified using an RNeasy RNA cleanup kit (Qiagen). Defolliculated stage V–VI *Xenopus laevis* oocytes were purchased commercially from Xenoocyte (Dexter, MI) and were stored in antibiotic supplemented ND96 (96 mM NaCl, 2 mM KCl, 1.8 mM CaCl_2_, 2 mM MgCl_2_, 100 units mL^−1^ penicillin, 100 μg mL^−1^ streptomycin) until use. Oocytes were microinjected with 2.5 ng cRNA (unless otherwise noted) and two-electrode voltage clamp (TEVC) recordings were performed 24 to 48 hours after microinjection.

For recordings, oocytes were impaled with borosilicate recording microelectrodes (0.3–2.0 MΩ resistance) backfilled with 3 M KCl. Oocytes were perfused with ND96 solution (96 mM NaCl, 2 mM KCl, 1.8 mM CaCl_2_, and 2.0 mM MgCl_2_, 5 mM HEPES [pH 7.4]) at a rate of 2.5 ml/min. Currents were evoked from a −90 mV holding potential by a 2000 ms ramp from −150 mV to +50 mV. Data were acquired using an OC-725C oocyte clamp amplifier (Warner Instruments) controlled by pClamp software (Molecular Devices), and digitized at 2 kHz using a Digidata 1332A digitizer (MDS Analytical Technologies).

For temperature experiments, recording solutions were heated by an SC-20 in-line heater/cooler combined with an LCS-1 liquid cooling system operated by the CL-100 bipolar temperature controller (Warner Instruments). Temperature was monitored using a CL-100-controlled thermistor placed in the bath solution immediately downstream of the oocyte. The perfusate was warmed from 15°C to 35°C in 5°C increments, with recordings performed once temperature readings stabilized at the desired values. For external pH experiments (pH_o_), each oocyte was initially perfused with ND96 solution at pH 7.4, switched to ND96 at pH 9.0 and then subsequently assayed at the experimental pH, with recordings performed 15 s after each new solution was applied. This allowed for all pH_o_ dependent changes to be normalized against the current density at pH 9.0. The buffer present in ND96 solutions was altered as appropriate, with one of the following: 10 mM TRIS (pH 9.0 - 8.1), 5 mM HEPES (pH 8.1-6.5) or 5 mM MES (pH 6.5 - 5.9). Aggregate pH_o_ data were fitted with a modified Hill equation: I = I_min_ + (I_max_ - I_min_)/(1 + 10^((pH_o_IC_50_/pH_o_) ∗ H)), where I_max_ and I_min_ are maximal and minimal current values, respectively, pH_o_IC50 is a half-maximal effective pH_o_ value, and H is the Hill coefficient.

For volatile anesthetic experiments, isoflurane saturated ND96 was prepared by adding commercially available isoflurane (Baxter Healthcare Corporation) to a 100ml volume of ND96 until phase separation was clearly observable. This mixture was stirred vigorously overnight in a closed glass bottle prior to use. The established maximal solubility of isoflurane in salt solution at room temperature is 15 mM (Scheller et al. 1997) and working isoflurane concentrations used for electrophysiological experiments were prepared by dilutions of the 15mM saturated stock solution. Azi-isoflurane was diluted into DMSO prior to use and the concentration of this stock solution was determined using the optical extinction coefficient of 126 M^−1^ cm^−1^ at 300nm Abs. Anesthetic solutions were stored in closed perfusion bags and were perfused onto oocytes through Teflon perfusion tubing, to prevent loss of isoflurane to the environment or electrophysiology rig.

## Results

### Azi-isoflurane photolabeling of TREK1 identifies VA binding sites

In order to identify TREK1 volatile anesthetic (VA) binding sites, we utilized a previously validated photo-reactive analog of the clinically important VA isoflurane (Eckenhoff et al. 2010). This biochemical adduct, azi-isoflurane, features a diazirine moiety capable of generating a highly reactive and chemically non-selective carbene adduct when irradiated with UV light. Azi-isoflurane has been shown to retain the anesthetic effects of isoflurane in animals and we first sought to ensure that the chemical modifications present in the azi-isoflurane compound would not alter the effect of this drug on the TREK1 channel. Two electrode voltage clamp studies of *Xenopus laevis* oocytes expressing TREK1 showed that application of azi-isoflurane causes a dose dependent potentiation of TREK1, with an EC_50_ of 735 ± 192 μM. We found no statistically significant difference between the effect of 2 mM isoflurane or 3 mM azi-isoflurane on TREK1 channel function (Fig. 1C).

**Figure 1:**
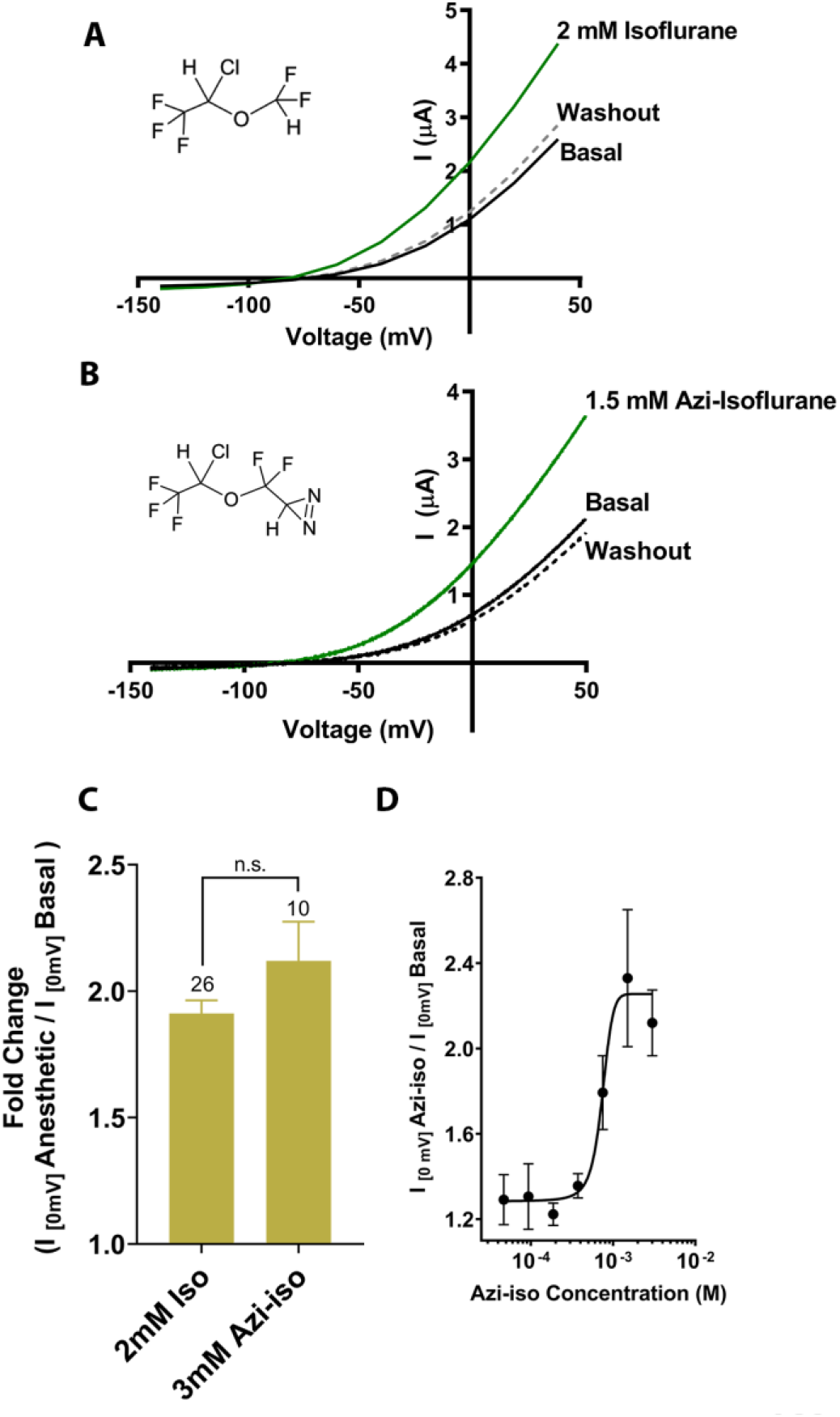
Functional validation of azi-isoflune activity on TREK1 channels. Representative two electrode voltage clamp recordings demonstrate the potentiating effects of saturating doses of isolflurane (A) and azi-isoflurane (B). Chemical structures of isoflurane and azi-isoflurane are shown. (C) Fold effect of administration of either isoflurane or azi-isoflurane on TREK1 outward current, as determined by the ratio of the recorded current at a voltage of 0mV, immediately prior to and following administration of VA agent. No significant difference was found between the responses of TREK1 to isoflurane versus azi-isoflurane, unpaired two tailed t-test P value of 0.11 (D) Dose response curve for azi-isoflurane activation of TREK1. Data derived from n>6, N>2 experimental observations. Error bars in panel C and D are mean ± SEM

For photolabeling studies, recombinantly expressed, purified and liposome reconstituted TREK1 protein (Supplemental Figure 1) was reacted with 30 μM azi-isoflurane, a concentration well below the EC_50_ of azi-isoflurane for TREK1, chosen to minimize non-specific modification. Following photolabeling, mass spectrometry (MS) of the TREK1 protein showed evidence of adduction of azi-isoflurane at two residues, G182 and K194, both located on the second transmembrane domain (TM2) of the channel (Figure 2). To examine whether the clinically relevant parent VA isoflurane also binds at these two sites, we performed a parallel azi-isoflurane photolabeling study of TREK1 in the presence of 3mM isoflurane as a competitive inhibitor. The 100-fold excess concentration of isoflurane protected the G182 site from azi-isoflurane photolabeling but did not prevent labeling at K194, suggesting that only the G182 site is specifically occupied by the parent VA isoflurane (Supplemental Figure 2).

**Figure 2:**
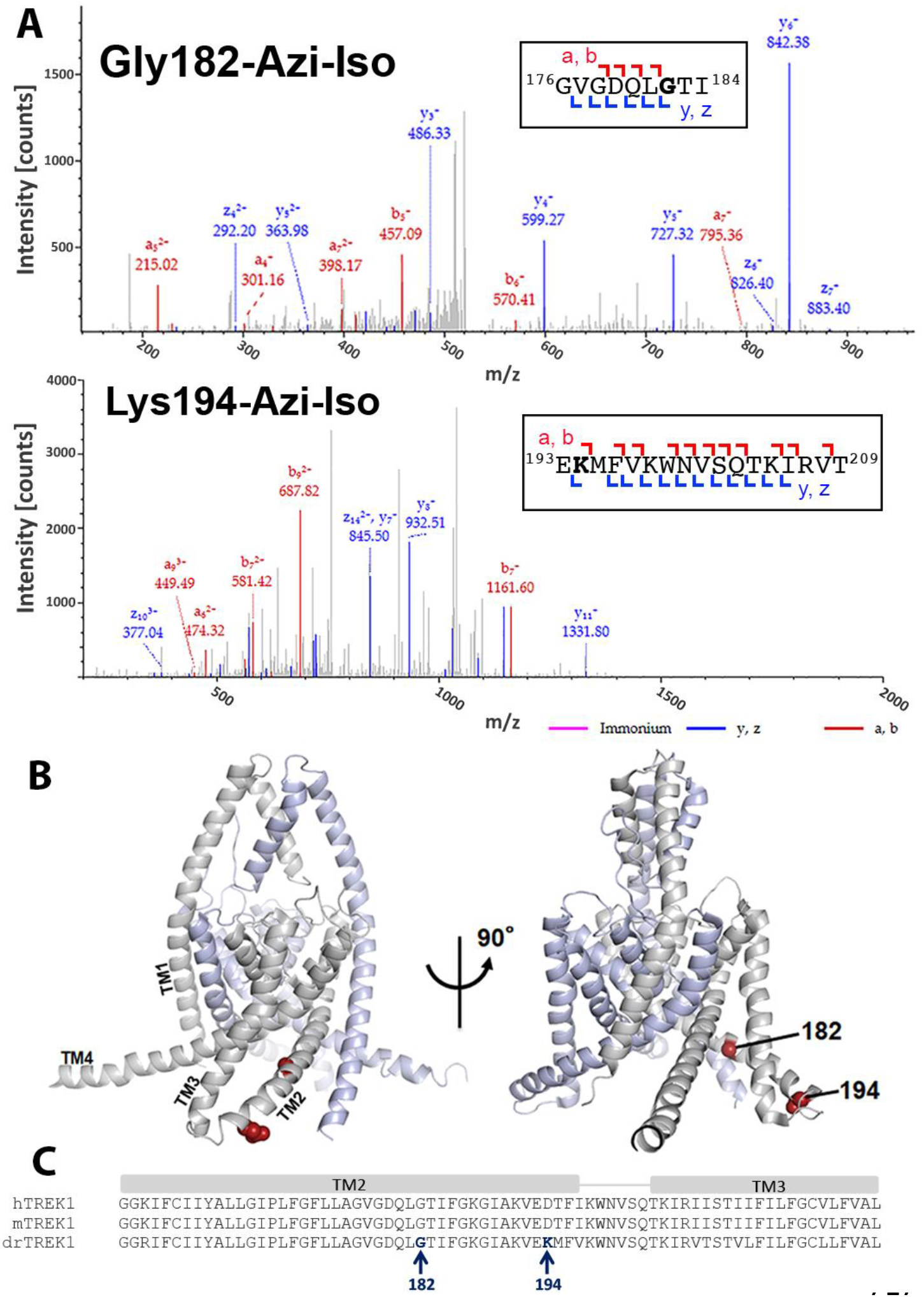
Azi-isoflurane photolabeling of TREK1: (A) Mass spectra of TREK1 photoaffinity labeled peptides labeled at Glycine182 (top) and Lysine 194 (bottom). Colored intensities denote the identified peptide a, b, z, and y ion fragments for the sequence assignment, as shown in the inset boxes. See Supp. Fig. 4 for corresponding peptide tables. (B) A structural model of mouse TREK1 (PDBID 6CQ6), showing the positions of residues G182 and K194 (labeled red spheres) along the TM2 helix. (C) Alignment of the TREK1 TM2 and TM3 helixes from human (hTREK1), mouse (mTREK1), and zebrafish (drTREK1).

While our MS results cover >91% of the TREK1 protein sequence with high confidence, there are notable regions of the TM3 and TM4 helices that are not identified by MS (Supplemental Figure 2). We attributed lack of coverage in these regions to the difficulty in ionizing and resolving hydrophobic transmembrane protein regions with this technique. Alternatively, binding of the hydrophobic azi-isoflurane photolabel could reduce the MS signal from bound protein peptides, causing gaps in MS coverage that correspond to regions where photolabeling has occurred. To exclude this possibility, we performed MS on non-photolabeled TREK1 protein, finding essentially the same lack of high confidence coverage within the TM3 and TM4 region. This suggests that the poor MS coverage of TREK1 TM3 and TM4 is not the result of photolabeling within these regions. However, we cannot rule out the possibility that azi-isoflurane might label positions within the TM3 or TM4 domains that we are simply unable to detect with our MS approach.

### Functional validation of TREK1 volatile anesthetic binding sites

To determine the functional importance of the VA binding sites identified by azi-isoflurane photolabeling, we introduced mutations at TREK1 positions 182 and 194 and used two-electrode voltage clamp recordings to assay for changes in TREK1 channel properties. We first substituted the endogenous amino acids at positions 182 and 194 with tryptophan, to mimic the size and hydrophobic nature of the azi-isoflurane photolabel. Tryptophan mutagenesis at position 194 (as well as other more conservative modifications) had no significant effect on the functional properties of the resultant mutant TREK1 channels (Figure 3). The amino acid at position 194 is poorly conserved across species, a lysine in the zebrafish TREK1 gene used for azi-isoflurane photolabeling but an aspartic acid in the mouse TREK1 construct used for our functional studies. To account for this difference, we introduced a D194K mutant into the mouse TREK1 background and found that this mutation also had no effect on TREK1 basal current or temperature dependence (Figure 3D, 3E). The absence of any observable functional effect of mutation at TREK1 194, along with the inability of excess isoflurane to protect position 194 from photolabeling by azi-isoflurane, suggest that despite being modified by azi-isoflurane, this site is unlikely to be relevant to the mechanism by which isoflurane and other clinically relevant VAs modulate TREK1. This notion is supported by the location of position 194 at the far end of TM2, facing toward the bulk solution and away from the core of the TREK1 protein.

**Figure 3:**
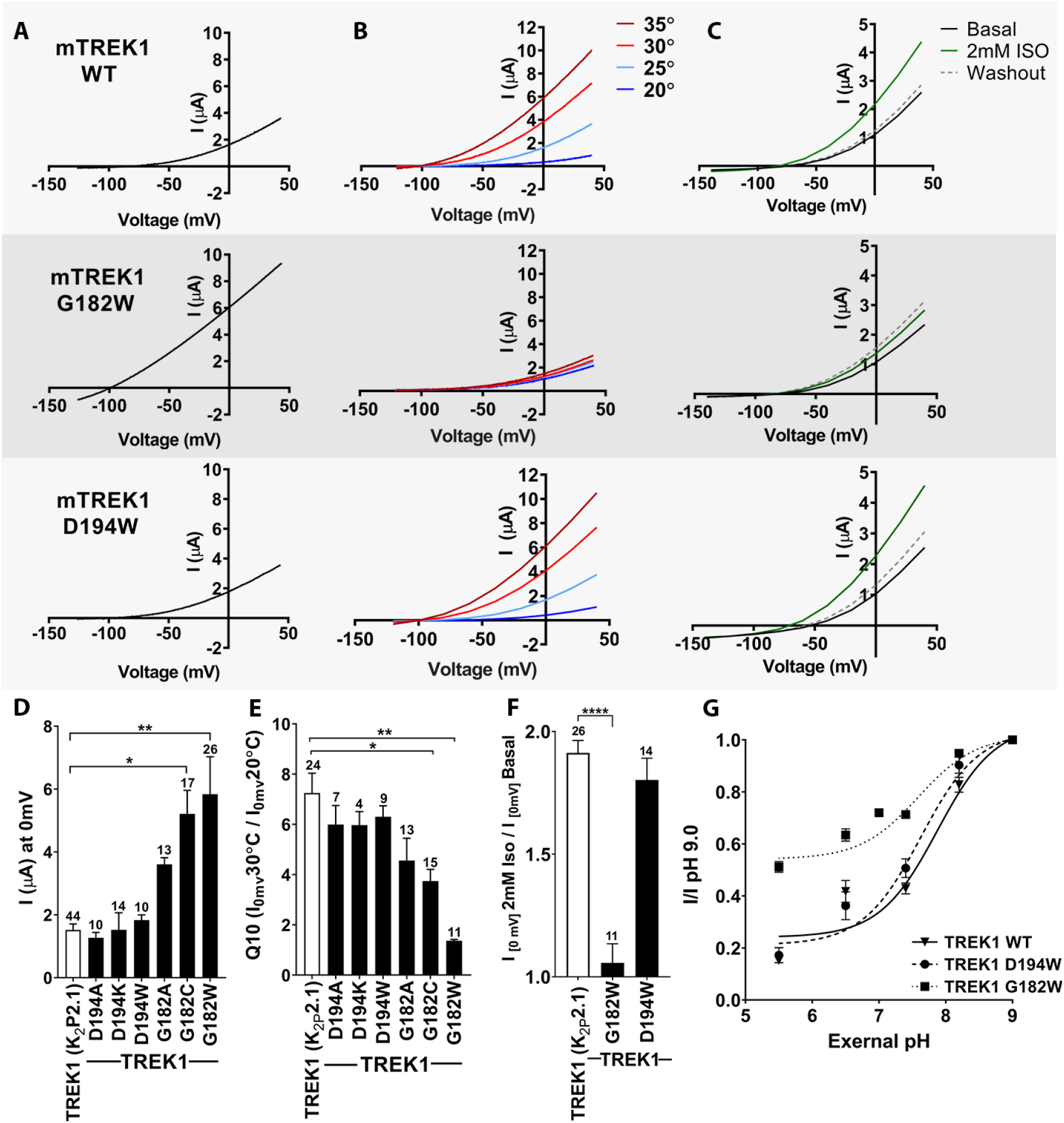
Function assessment of TREK1 residues identified by azi-isoflurane photolabeling: Representative two electrode voltage clamp recordings of TREK1 wildtype and mutant channels TREK1 G182W and D194W. (A) Basal currents measured 24 hours after microinjection of 2.4 ng cRNA. (B) Temperature dependence of TREK1 currents, measured at temperatures of 20°C to 35°C, in 5°C increments. (C) Response to administration of 2mM Isoflurane, followed by washout. For temperature and isoflurane experiments performed on TREK1 channels bearing mutations that alter basal current density, the concentration of microinjected cRNA was titrated to achieve 1 uA of current at 20°C, to approximate wildtype channel current density. (D) Quantification of TREK1 channel activity on basal current level, (E) temperature dependence as measured by Q10 (30°C/20°C), (F) response to isoflurane administration, or (G) changes in external pH, as measured by TREK1 current at 0 mV. Number of replicate experiments indicated. Error bars are mean ± SEM. Statistically significant results indicated, * p<0.5, ** p<0.05, **** p<0.0005.

TREK1 G182 is located within the center of TM2, one helical turn away from a glycine at position 178 previously shown to be a hinge point for a buckling motion of the TM2 helix that occurs during K2P gating (Lolicato et al. 2014). Tryptophan mutagenesis at TREK1 G182 demonstrated a large increase in basal current density and a near complete loss of modulation by heat or isoflurane, findings suggestive of TREK1 channel activation (Figure 3A-C, note that in panel 3B and 3C the concentration of injected cRNA was adjusted to normalize basal current density between mutant and WT TREK1 channels). Given the unique ability of glycine residues to impart helical flexibility and the known conformational movements in the region of TREK1 around G182 (Lolicato et al. 2014, Brohawn et al. 2014a, Dong et al. 2015), we sought to determine whether loss of flexibility at position 182 was responsible for the major functional effects observed in the TREK1 G182W mutant. By introducing more conservative mutations at G182 we discovered that the effect of mutagenesis best correlated with the size of the introduced amino acid. TREK1 G182A showed only a small potentiating effect on current density and no significant effect on gating by heat and TREK1 G182C showed an intermediate phenotype (Figure 3D-F).

To further demonstrate the potentiating effect of increased side chain size at TREK1 G182, we treated TREK1 G182C channels with MMTS, a membrane permeant cysteine-modifying reagent (Figure 4). In wild type TREK1 channels, MMTS treatment caused an increase in channel activity that washed out within 2-3 minutes of removal of MMTS, a seemingly non-specific effect that occurred even in a TREK1 construct with all five endogenous cysteines removed (TREK1 cys-). By contrast, modification of the TREK1 G182C mutant with MMTS caused potentiation of channel activity that persisted after washout of the MMTS reagent (Fig. 4A). An equivalent effect was observed in a TREK1 cys-G182C mutant, specifically confirming that it is modification of the G182 cysteine that leads to the observed persistent potentiation of TREK1 currents following MMTS application (Fig. 4B). These findings support the notion that steric crowding is a significant determinant of the potentiation observed after perturbation of the G182 residue, whether by mutagenesis, MTS reagent modification, or by occupancy with a VA agent or photolabel.

**Figure 4:**
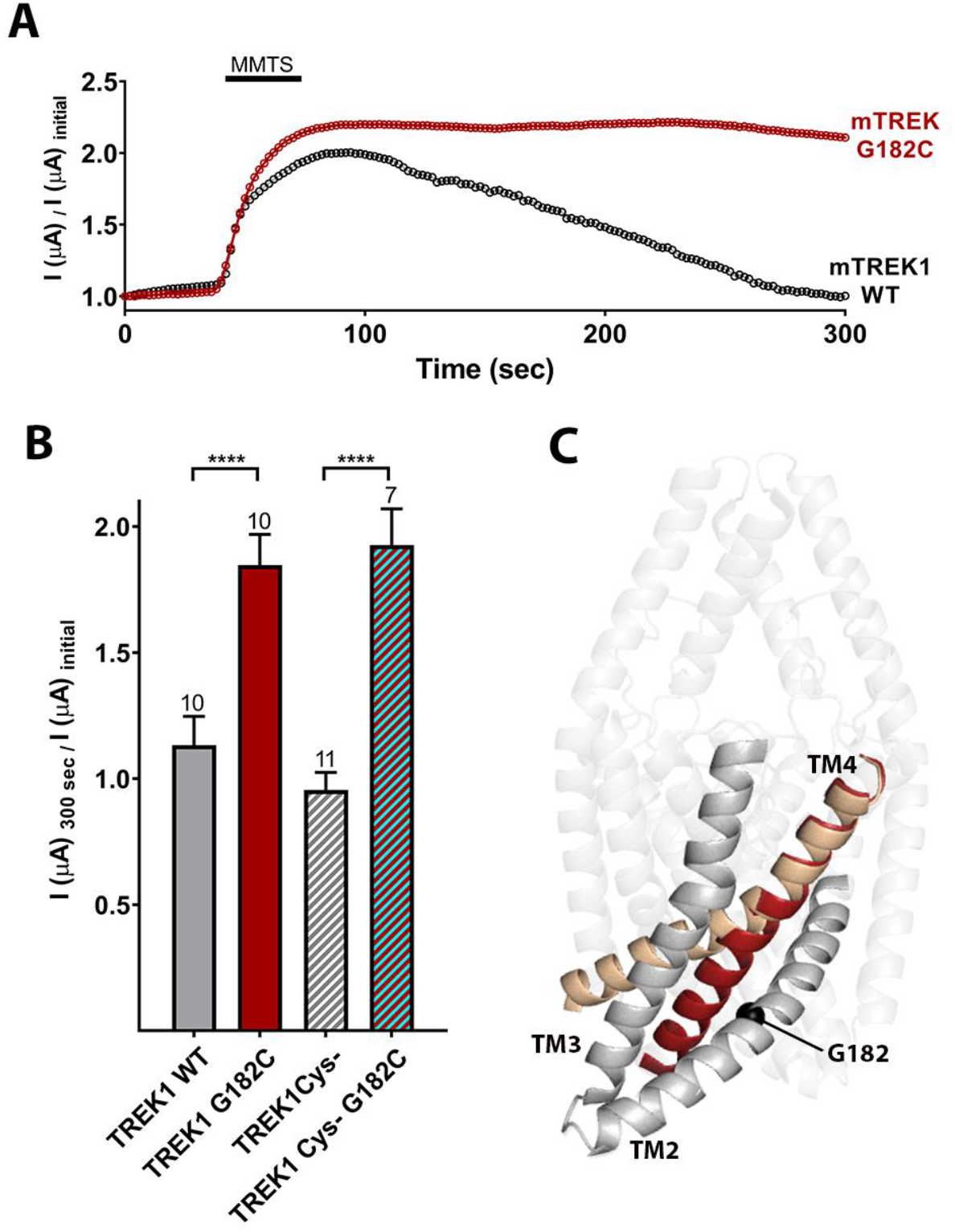
Cysteine modification at the G182 position leads to TREK1 activation: (A) Representative time courses of the response of TREK1 WT or TREK1 G182C channels to treatment with 2mM MMTS. Shown is the recorded current at 0mV holding potential, measured every 5 seconds for 5 minutes, with current values normalized to the initial value recorded at the beginning of the time course (B) Quantification of effect size at the end of the time course (300 seconds), for TREK1 WT and TREK1 Cys-, a mutant TREK1 channel lacking all 5 of the endogenous TREK1 cysteine residues, and for G182 mutants in both of the above backgrounds. Number of replicate experiments indicated. Error bars are mean ± SEM. Statistically significant results indicated, **** p<0.0005. (C) Crystallographically defined structural models of TREK2 in the TM4 up (PDB ID 4xdl, tan) and TM4 down (PDB ID 4bw5, maroon) conformations (Dong et al. 2015), highlighting the TM2, TM3, and TM4 helices from only a single subunit. The position of G182 (black sphere) is noted.

To explore whether disruption of isoflurane binding contributes to the VA insensitivity of the G182W mutant channel, we performed azi-isoflurane photolabling studies on purified TREK1 G182W protein. We found no evidence of azi-isoflurane labeling at position 182 or at any neighboring residues, suggesting that the G182W mutation either eliminates azi-isoflurane binding or reduces the affinity of VA for this site. We did observe azi-isoflurane photolabeling at two positions far from the G182 site (Supplemental Figure 2), one near the top of TM1 (A67) and one in the C-terminal domain (T303). These residues were not photolabeled in the VA sensitive TREK1 WT background and as such they are unlikely to contribute to the mechanism by which VAs modulate TREK1. However, enhanced accessibility of these residues in only the G182W background could be explained by global changes in protein tertiary structure induced by the G182W mutation, consistent with the dramatic functional effects observed in this mutant.

While the G182 residue is located at the cytosolic face of the TREK1 channel, activation gating in K2P channels is thought to occur via a “C-type” gating mechanism involving a rearrangement of the extracellular selectivity filter region (Bagriantsev et al. 2011, Piechotta et al. 2011, Zilberberg, Ilan and Goldstein 2001). Given that the G182 residue is located far from the selectivity filter (Figure 2B), a significant allosteric effect would be required for steric crowding at G182 to exert an effect on this distant gate. To explore whether the G182W mutation modulates the TREK1 “C-type” gate, we examined the stability of the selectivity filter against closure by application of extracellular acidosis. Acidification of the extracellular face of TREK1 has been shown to inhibit TREK1 channel activity and to simultaneously diminish potassium selectivity, a hallmark of selectivity filter based gating (Cohen et al. 2008, Sandoz et al. 2009). While the TREK1 WT and TREK1 D194W mutants were both strongly inhibited by application of extracellular acidosis, the TREK1 G182W channel was resistant to closure by acid (Figure 3G, IpH_5.5_/pH_9.0_ 0.15 ± 0.05 for TREK1 WT, 0.17 ± 0.1 for TREK1 D194W, 0.51 ± 0.08 for TREK1 G182W), though the IC_50_ of the effect was similar for all three channels tested (pH 7.88 ± 0.07 for TREK1 WT, 7.63 ± 0.08 for TREK1 D194W, and 7.59 ± 0.07 for TREK1 G182W). These data support a mechanistic model in which isoflurane modulation of the TREK1 channel occurs via anesthetic binding to the G182 site causing allosteric stabilization of the selectivity filter “C-type” gate, which leads to increased TREK1 channel activity.

Many of the biophysical modalities known to modulate TREK1 channels, including heat, mechanical stretch, intracellular acidosis, and bioactive lipids, are believed to alter channel activity by affecting the selectivity filter “C-type” gate (Bagriantsev et al. 2011, Bagriantsev, Clark and Minor 2012, Piechotta et al. 2011), though the input sensor for these biophysical gating cues was proposed to be the intracellular C-terminal domain (Honore et al. 2002, Chemin et al. 2005, Bagriantsev et al. 2012). Structurally defined “TM4 up” and “TM4 down” conformational states (Brohawn et al. 2014a, Lolicato et al. 2014, Dong et al. 2015) have been identified as the key rearrangements that allow these C-terminal domain inputs to traverse the protein and converge at the selectivity filter to modulate gating (Zhuo et al. 2016). Within this gating model, the location of the G182 anesthetic binding site along the TM2 helix in direct opposition to TM4 (Figure 4C) is intriguing. It suggests that VA occupancy at the G182 site could modulate TREK1 activity by influencing TM4 intra-molecular rearrangements known to play a key role in K2P gating. To further explore the isoflurane binding site and gain insight into the effect of anesthetic binding on TREK1 channel gating, we utilized molecular dynamics (MD) simulation.

### MD simulation identifies residues important for VA modulation to TREK1 channels

In order to establish a suitable starting point for MD simulations, we evaluated potential ligand binding configurations compatible with our photoaffinity results, using Autodock Vina docking software (Trott and Olson 2010). A pocket formed by G182 and the nearby TM3 and TM4 helices had the highest ranked score of −5.1, and was the only predicted binding site near the G182 residue. This binding site is located entirely within the transmembrane region of TREK1 and isoflurane is a relatively low-affinity ligand with minimal electrostatic interactions. Our docking results confirmed that the binding site near G182 was a sterically favorable starting point to place isoflurane for subsequent equilibrium MD simulation. Simulations of TREK1 embedded in a POPC:cholesterol lipid bilayer were conducted of TREK1 wildtype (440 ns) and TREK1 G182W in the absence of isoflurane (960 ns), and of TREK1 wildtype (WT) with a single isoflurane molecule placed at the G182 site in one of the two K2P subunits (700 ns). A representative snapshot of the isoflurane binding site is depicted in Figure 5A.

**Figure 5:**
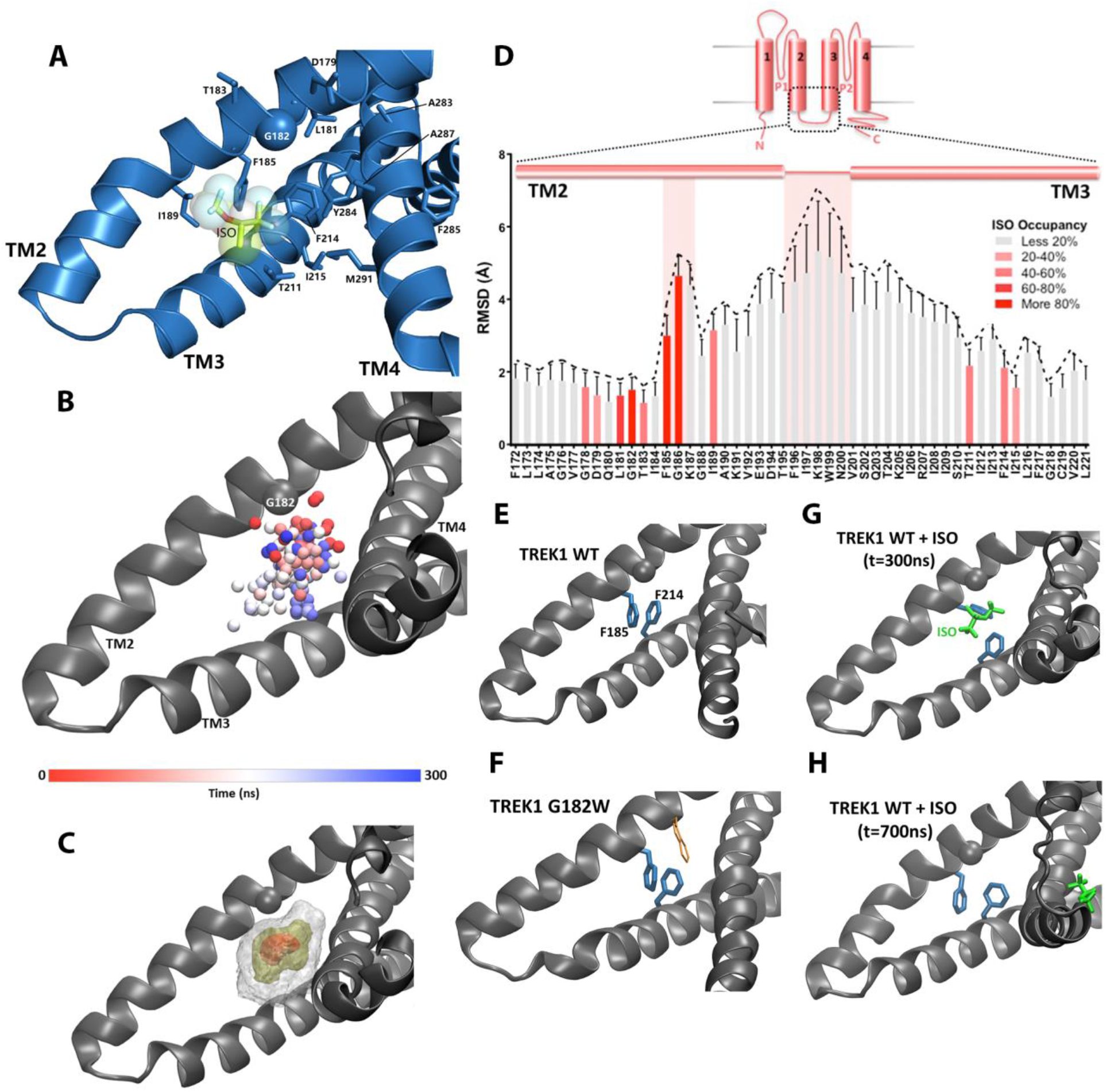
A) Isoflurane binding pocket predicted by MD simulation. B) Position of the oxygen atom at the center of the isoflurane molecule during the first 300ns of equilibrium MD simulation of TREK1 WT in the presence of isoflurane C) Density map of the position of the bound isoflurane. Isosurfaces represent 10%, 30%, and 50% isoflurane occupancy. D) Mean RMSD of residues within the TREK1 TM2/TM3 loop during the ~360 ns when isoflurane was bound, as compared with equilibrated free wild-type TREK1. Relative isoflurane occupancy for each residue (see Table 1) is shown as quartiles, described on right. Regions demonstrating the largest RMSD deviations are highlighted. E) Representative snapshot of equilibrium MD simulations of TREK1 WT or F) G182W show pi-stacking of the F185 and F214 residues. Equilibrium MD simulations of TREK1 in the presence of isoflurane reveal disruption of F185/F214 pi-stacking, both during the period that isoflurane remains in the binding pocket (panel G, t=300ns) as well as after isoflurane has left the binding pocket (panel H, t=700ns)

Simulation of WT TREK1 in the presence of VA demonstrated isoflurane to be highly mobile within the binding pocket, adopting a range of positions (figure 5B) and ultimately escaping the binding site after approximately 360 ns of simulation. This observed high degree of isoflurane mobility is consistent with both the low binding affinity predicted by docking analysis and the high micromolar concentrations of isoflurane required to activate TREK1 in functional assays. Despite the mobility of the isoflurane ligand, we were able to identify a number of residues that show greater than 20% occupancy by isoflurane (Table 1, Figure 5A), with occupancy defined as the percentage of MD trajectory snapshots (prior to isoflurane escaping the binding site) in which isoflurane was located within 7 Å of a given residue. High occupancy positions (Figure 5A) include the G182 residue and additional amino acids on TREK1 TM2, TM3, and TM4, as listed in Table 1. In agreement with our photolabeling findings, MD simulations demonstrated that isoflurane remained in close proximity to the G182 residue for >94% of the time VA occupied the binding site. This finding supports the notion that isoflurane imposes a consistent steric crowding at the G182 position, akin to the activity enhancing modifications we introduced through either mutagenesis or biochemical modification.

**Table 1.** Occupancy of residues in G182 isoflurane binding pocket, defined as the percentage of snapshots where isoflurane within 7 Å of the given residue during MD simulation. Residues with occupancy less than 20% are omitted. Positions previously shown to mediate VA sensitivity in TASK K2P channels are annotated (*)

| Mouse TREK1<br>Residue Number | Percent Occupancy<br>During MD Simulation | Transmembrane Domain |
| --- | --- | --- |
| GLY178 | 47.65% | TM2 |
| ASP179 | 35.19% | TM2 |
| LEU181 | 68.40% | TM2 |
| GLY182 | 94.01% | TM2 |
| THR183 | 69.07% | TM2 |
| PHE185* | 92.17% | TM2 |
| GLY186 | 89.16% | TM2 |
| ILE189 | 52.96% | TM2 |
| THR211* | 56.20% | TM3 |
| PHE214 | 44.39% | TM3 |
| ILE215 | 25.65% | TM3 |
| ALA283 | 28.07% | TM4 |
| TYR284 | 67.95% | TM4 |
| PHE285 | 27.41% | TM4 |
| ALA287 | 88.05% | TM4 |
| VAL288 | 78.20% | TM4 |
| MET291* | 56.11% | TM4 |

The proximity of the VA binding site to the neighboring TM4 helix (Figure 4C) suggested a mechanism by which anesthetic binding could influence K2P activity by influencing TM4 position, which is bolstered by our MD simulation data. We found a number of TM4 residues that directly interact with the bound isoflurane molecule, most notably Y284 and M291. TREK1 Y284, the only polar residue in the otherwise hydrophobic TM4 helix, has been proposed to form stabilizing hydrogen bonds with one of two alternative TM3 backbone carbonyls in the “up” versus “down” conformations of TM4 (Dong et al. 2015), but is highly occupied by isoflurane when the drug is bound in the VA site. Similarly, the M291 position has been proposed to form TM4 conformation dependent interactions with neighboring TM3 and TM4 residues and in our simulation M291 is 55% occupied by the bound isoflurane molecule.

To assess for structural changes in the TREK1 protein induced by the presence of bound isoflurane, we compared the MD simulations of isoflurane-free versus isoflurane-bound TREK1 and calculated the per residue root mean square deviation (RMSD) of the TREK1 protein during the two simulations. The isoflurane molecule did not induce any major conformational changes in TREK1 during our simulations and we did not observe any alteration in the position of the TM4 helix attributable to the presence of isoflurane. However, within the VA binding pocket, we found an increase in RMSD along a short span of the TM2 helix (Figure 5C) that contains residues exhibiting the highest measured isoflurane occupancy. This localized structural change induced by the presence of isoflurane alters the positioning of the F185 residue, a residue previously shown to impact K2P responsiveness to heat, volatile anesthetics, mechanical stretch, and the pharmacological efficacy of BL1249, a K2P activator (Lolicato et al. 2014, Dong et al. 2015, Luethy et al. 2017, Pope et al. 2018). The aromatic group of the F185 residue is known to form a pi stacking interaction with the TM3 F214 residue, an interaction proposed to stabilize the “TM4-down” gating conformation in TREK2 channels (Dong et al. 2015).

To explore the effect of isoflurane binding on this structurally important interaction, we quantified pi-stacking between F185 and F214 in our simulations, such that the two aromatic rings were considered to be pi-stacked if their centroids were separated by 4.4 Å or less and the angles between the ring planes were less than 30 degrees (McGaughey, Gagne and Rappe 1998). In the TREK1 WT and TREK1 G182W simulations, pi-stacking between these residues occurred in 37% and 38% of trajectory frames, respectively, but pi-stacking occurred in only 2% of the trajectory frames in the isoflurane-bound wildtype TREK1 simulation. Disruption of this interaction was caused in large part by the presence of the isoflurane molecule itself distorting the binding pocket (Figure 5C, 5D), though we found that pi stacking between F185 and F214 remained disrupted for an additional ~300 ns of MD simulation time even after the isoflurane molecule had left the VA binding site. We surmise that this persistent disruption of the F185/F214 interaction is due to the observed local deformation of the TM2/TM3/TM4 interface induced by the presence of isoflurane, as shown in Figure 5D.

Based on these findings, we hypothesize that a combination of VA induced local structural perturbations, disruption of the F185/F214 pi stacking interaction, and direct steric effects upon TM4 movement all combine to alter the dynamics of the TM4 movements known to influence K2P gating. While our MD simulations did not directly reveal major conformational rearrangements of TM4, the observed isoflurane occupancy at numerous positions known to govern the energetics of TM4 translocation from the “down” to “up” position suggest a mechanism by which VA binding ultimately alters K2P activity by modulating TM4 movements.

### Key residues transfer VA insensitivity from TRAAK to TREK1

While both TREK and TASK channels are potentiated by VA agents, the TRAAK K2P channel is anesthetic insensitive (Patel et al. 1999). TRAAK is 44% identical and 69% homologous to TREK1 and the two channels share many functional properties, including modulation by heat, mechanical stretch, pH, and arachidonic acid (Maingret et al. 1999, Kang, Choe and Kim 2005). The relative selectivity of VA agents for TREK1 over TRAAK despite the many similarities between these two channels allowed us to utilize the TRAAK channel as a negative control to further characterize the K2P VA binding site.

VA insensitivity of TRAAK could arise from one of two possibilities; failure of VAs to bind to TRAAK or inability of bound VA to modulate TRAAK activity. To distinguish between these possibilities and ascertain whether VAs bind to the TRAAK channel, we recombinantly expressed, purified and reconstituted TRAAK into liposomes (Supplemental Figure 1) and performed azi-isoflurane photolabeling studies identical to those used for TREK1 channels. The G182 residue identified by azi-isoflurane photolabeling of TREK1 is conserved in the TRAAK sequence, but we found no evidence of azi-isoflurane photolabeling at this glycine or at any other position in the TRAAK channel. MS of TRAAK protein after photolabeling showed a relatively lower total coverage of the TRAAK sequence compared to our results for TREK1 (84% for TRAAK versus 91% for TREK1), but all TRAAK residues homologous to the TREK1 azi-isoflurane binding region identified by MD simulations were positively identified, precluding the possibility that azi-isoflurane binds within this region of TRAAK. Our results suggest that the inability of VA agents to activate TRAAK channels is due to either an absence of binding or a significantly reduced affinity of anesthetic for this VA binding region.

We next explored the sequence similarity between TREK1 and TRAAK at the residues that showed the highest isoflurane occupancy during MD simulations. Amongst the identified amino acids, we found four residues that differ significantly between the TREK1 and TRAAK sequences (Figure 6A-B), and created a series of mutant TREK1 channels that contain one or two of the corresponding TRAAK residues at these positions (Fig. 6C-D). When examined by TEVC, the “TRAAK-like” mutant TREK1 channels all exhibit diminished responsiveness to the potentiating effect of isoflurane (Figure 6C), supporting a role for these residues in either forming the VA binding domain or transducing the effects of binding to channel opening. While the extent of the effect of these mutants was less dramatic than the near complete absence of isoflurane responsiveness in the G182W mutant, the effect of combining two TRAAK-like mutations was additive, as shown for the double mutant TREK1 F185L G186R (Figure 6C).

**Figure 6:**
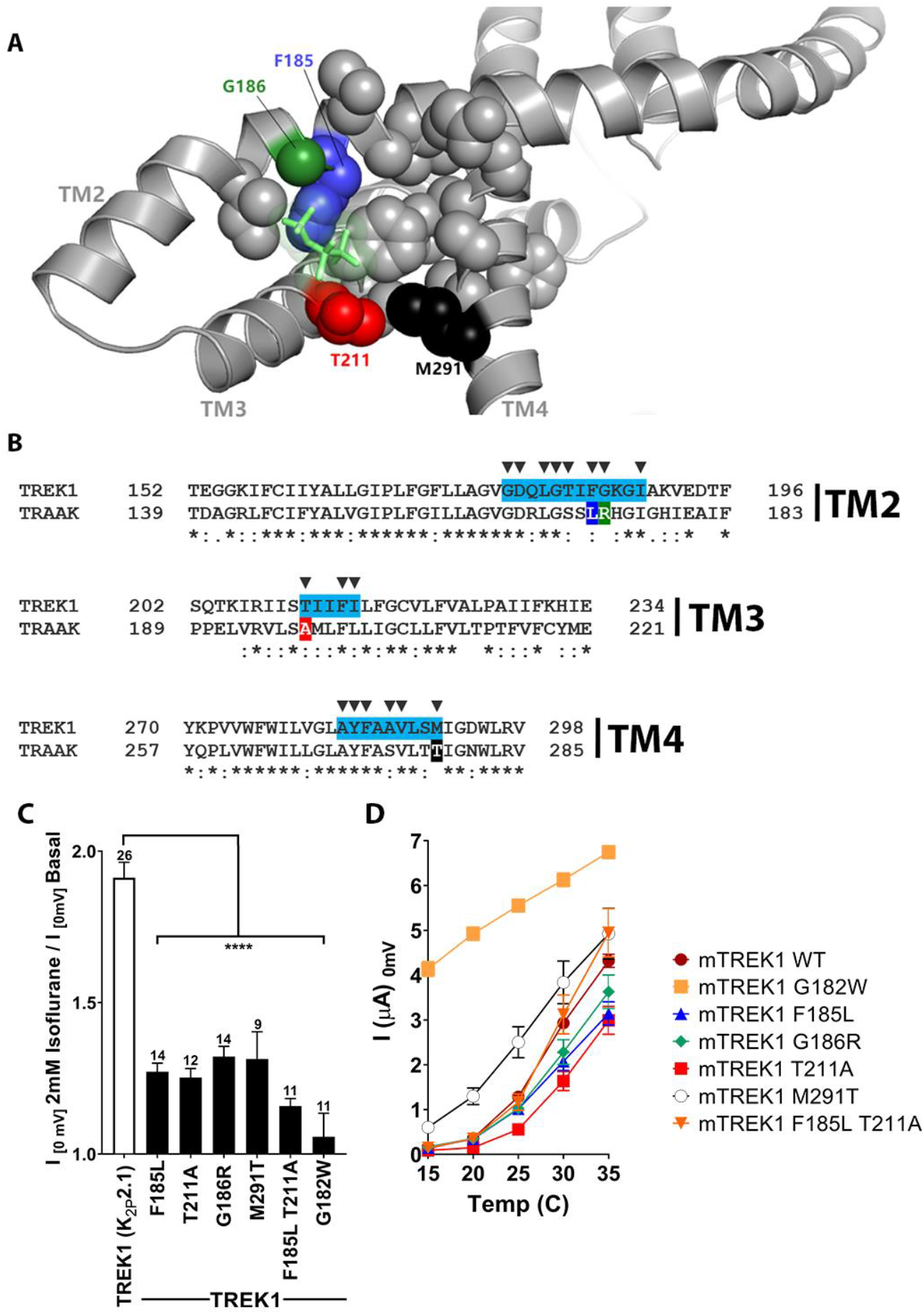
Mutations in the isoflurane binding site alter anesthetic sensitivity without perturbing global channel function. (A) Representative MD snapshot showing the isoflurane binding site, including residues predicted to have >20% occupancy by isoflurane shown in sphere representation, isoflurane show in stick representation (green). (B) Alignments of mouse TREK1 and TRAAK sequences, with the isoflurane binding domain regions of TREK1 TM2, TM3, and TM4 (as identified by molecular dynamics simulation) highlighted in blue. Arrows denote positions of high isoflurane occupancy. Poorly conserved residues are color coded throughout the figure [F185 (blue), G186 (green), T211 (red), M291 (black)]. (C) Quantification of TREK1 wildtype and mutant responses to 2mM isoflurane administration or (D) temperature, as measured by TREK1 current at 0 mV. Number of replicate experiments indicated. Error bars are mean ± SEM. Statistically significant results indicated, **** p<0.0005.

Whereas the G182W mutant exhibited diminished responsiveness to both VA activation and heat, the “TRAAK-like” mutants maintain a preserved sensitivity to activation by heat (Figure 6D). The fact that the “TRAAK-like” mutants selectively alter VA activation without perturbing gating by heat argues for their specificity to the mechanism of action of VAs. Activation of TREK1 by heat is known to be critically related to the phosphorylation state of serine 333 in the intracellular C-terminal domain of TREK1 (Maingret et al. 2000) and the persistence of TREK1 heat activation in these “TRAAK-like” mutants suggests that these mutations do not eliminate VA modulation by simply causing a global activation of the TREK1 “C-type” gate (as was the case for the G182W mutant). Rather, the ‘TRAAK-like” mutations appear to reduce VA sensitivity by acting specifically at the VA site of action, consistent with predictions made by our MD simulation data and TREK1 vs TRAAK sequence homology.

## Discussion

Effort to determine regions within the K2P channel structure susceptible to small molecule modulators has become a topic of significant recent interest (Dong et al. 2015, Lolicato et al. 2017, Pope et al. 2018, Rinné et al. 2019), motivated by the lack of specific and high affinity pharmacology targeting K2P channels. Here we focus on K2P modulation by VAs, drugs heavily utilized in common clinical practice. We identify an interface formed by the TM2, TM3, and TM4 helices of TREK1 as the binding site for the VA isoflurane and identify site-specific interactions between TREK1 and isoflurane that define this binding region. The identified VA modulatory site features TREK1 residues previously identified to play key roles in K2P modulation by mechanical stretch, heat, and the pharmacological activator BL1249, suggesting a common mechanistic pathway shared by VA and other K2P gating cues.

Prior to our study, the strongest evidence for a VA binding site in K2P channels was found in the TASK subfamily, where a single point mutation in the *lymneal* TASK TM3 residue M159 (homologous to TREK1 position T211) caused loss of steroselective discrimination between R and S optical isomers of isoflurane (Andres-Enguix et al. 2007). TREK1 T211 lies within a span of the TM3 helix we were unable to resolve with MS but through MD simulation we identified T211 as a determinant of TREK1 VA binding (Table 1). Two additional residues identified by MD simulation, TREK1 F185 and M291, also align to positions implicated in TASK VA sensitivity (Andres-Enguix et al. 2007, Conway and Cotten 2012). All three of these residues (F185, T211, M291) differ significantly in the TRAAK protein sequence and “TRAAK-like” mutations at these positions decreased TREK1 VA sensitivity (Figure 6B). The overlapping molecular determinants of TREK1 and TASK1 VA sensitivity (and of TRAAK VA insensitivity) suggest a shared VA binding site across the K2P family. While the gating mechanisms of TASK channels have not been as extensively studied as those of the mechanosensitive K2Ps, recent work exploring the inhibitory effect of the local anesthetic agent bupivacaine on TASK1 channels demonstrates that bupivacaine binding constrains TM4 helix movement and ultimately alters selectivity filter behavior (Rinne et al. 2019). The bupivacaine site of action in TASK channels differs from the VA modulatory site identified in our study, but the effect of drug binding on TM4 conformation may be a shared feature in K2P gating utilized by both local and general anesthetics.

Our data suggest that VAs modulate K2P channel activity by influencing the position of the TM4 helix. The proximity of the azi-isoflurane labeled TREK1 G182 residue to the TM4 helix initially suggested such a mechanism and the potentiating effect of increased G182 side chain size supported this hypothesis. MD simulation studies identified direct contacts between isoflurane and the TM4 helix. However, the MD simulation data do not show major TM4 conformational rearrangements in the presence of isoflurane and thus whether VAs induced TM4 movements. Similarly, we have no evidence of whether the VA bound TREK1 channel more closely resembles the “TM4 up” or “TM4 down” states defined by crystallography (Dong et al. 2015, Brohawn et al. 2014a). As there is evidence to suggest that both the “TM4 up” and “TM4 down” conformations can promote active K2P channel states (McClenaghan et al. 2016), each of the two conformations could be consistent with a VA activated channel. However, both the direct steric constraints imposed by the presence of VA near TM4 and the disruption of the “TM4 down” state stabilizing F185/F214 pi-stack interaction would be predicted to favor a “TM4 up” conformation. The starting point for our MD simulations is a crystallographically-determined structural model of TREK1 already in the “TM4 up” state (Lolicato et al. 2017) and may explain the lack of any additional TM4 rearrangement in the presence of isoflurane. Further MD simulation utilizing a homology model of TREK1 based on K2P structures determined in the “TM4 down” state might more definitively answer this question. However, it is also possible that VA binding induces a conformation distinct from either of these two structurally defined states. The molecular mechanisms governing the connectivity between TM4 conformation and the flux gating behavior of the selectivity filter (Schewe et al. 2016) remains poorly defined and is an area of significant ongoing interest.

Having identified a VA modulatory site that appears to be shared across the TREK and TASK K2P subfamilies, we are cognizant of some of the inconsistent VA responses observed across these K2P subfamilies. For instance, while the obsolete VAs chloroform and diethyl-ether activate TREK1 and TASK3, they paradoxically inhibit the TASK1 channel (Patel et al. 1999, Andres-Enguix et al. 2007). Nitrous oxide and xenon have been reported to activate TREK1 but have no effect on TASK channels (Gruss et al. 2004), and TRAAK channels are unresponsive to VAs despite their close similarity to other mechanosensitive and VA sensitive K2Ps. Meanwhile, little is known about the gating of THIK channels but they are inhibited by the VAs halothane and isoflurane, an effect that appears to play an important role in the clinically relevant central respiratory depressant effect of VA administration (Rajan et al. 2001, Lazarenko et al. 2010). How can we explain the paradoxical effects of VA agents at structurally similar targets? One explanation would be that the VA binding site identified in our study represents only one of a subset of VA modulatory domains in the K2P family. Within the anesthetic mechanisms literature there are numerous examples of VA responsive ion channels that feature multiple low affinity anesthetic modulatory sites acting in a combinatorial fashion to produce a given output (Fourati et al. 2018, Heusser et al. 2018, Woll et al. 2018, Hemmings et al. 2019). However, the location of the identified K2P VA binding site at a position central to TM4 gating offers an alternative explanation. We understand relatively little about the mechanism by which TM4 movements govern K2P channel activity and it is possible that VA binding to this single site may produce opposing effects in the context of subtle differences in the connectivity between TM4 movements and the channel gate. Recent studies utilizing the state dependent binding of the mechanosensitive K2P inhibitor fluoxetine demonstrate that the resting and activated conformation of the TM4 helix in TREK1 versus TRAAK channels are inverted, underscoring this point (Soussia et al. 2018) and perhaps contributing to the absence of VA binding and responsiveness in TRAAK channels.

While our results support the presence of a defined modulatory site within the TREK1 channel structure, these findings do not preclude a contribution from indirect effects of VAs on membrane lipid composition, as has been recently proposed (Pavel et al. 2019). Such a mechanism is in fact a particularly appealing explanation for the effects of the structurally simple and TREK1 specific VA agents chloroform, nitrous oxide, and xenon (Gruss et al. 2004). In this context, the location of the TREK1 VA binding site described in our study at a region key to the mechanism by which C-terminal domain dependent signals (including lipid modulation) are translated to TM4 (Zhuo et al. 2016) and ultimately to the selectivity filter gate would allow both an indirect and direct modulatory effect on channel behavior to feed into a shared common pathway.

---

**Supplemental Figure 1:**
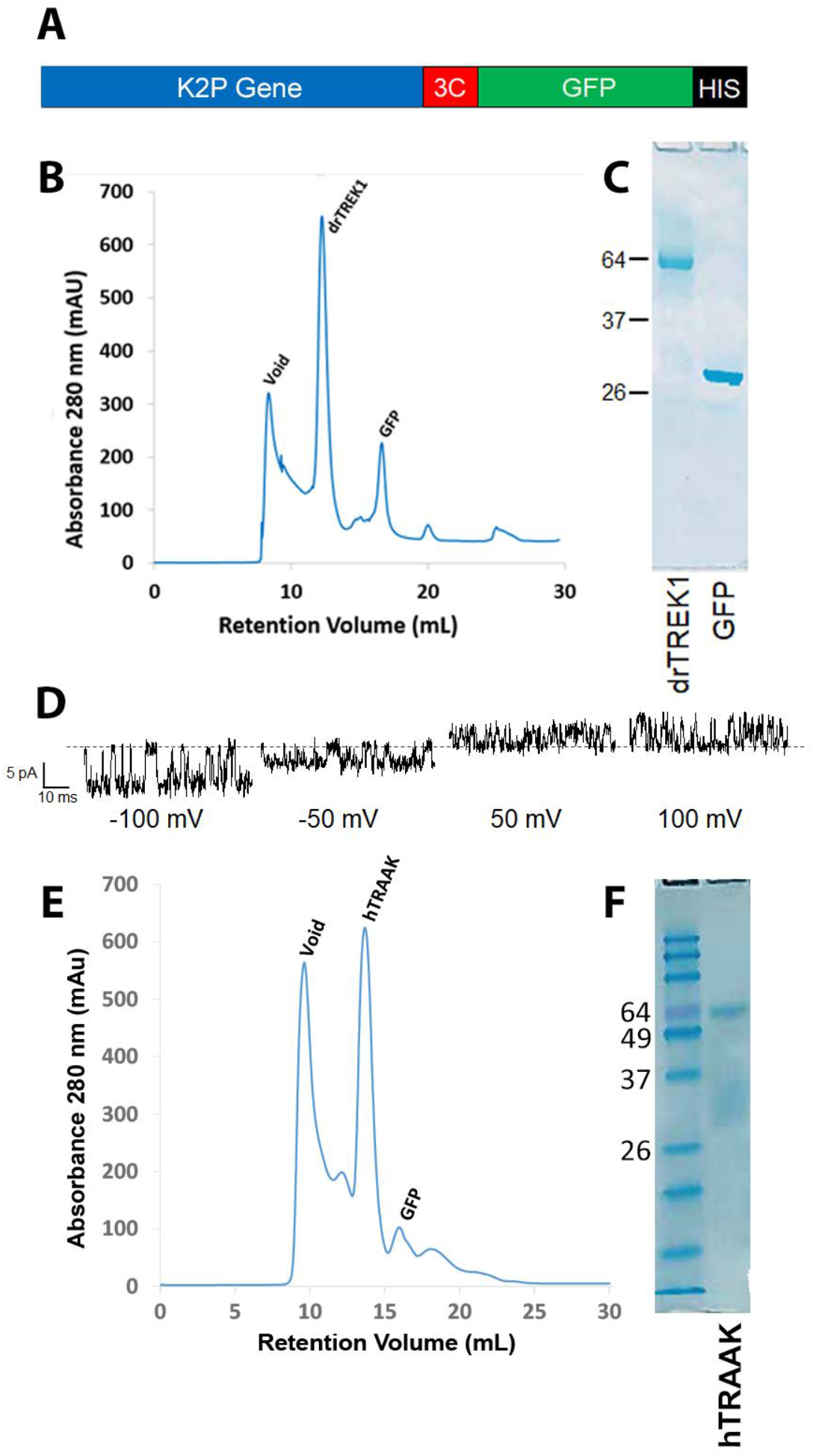
Purification of TREK1 and TRAAK proteins. (A) Zebrafish TREK1 or human TRAAK DNA sequences were purified by metal chromatography via a C-terminal HIS tag, and expressed as fusion proteins with GFP, cleaved off via a 3C protease cleavage site prior to (B,E) final size exclusion chromatography. (C, F) After SDS PAGE electrophoresis, purified protein ran at a molecular weight of approximately 65 kDa, consistent with a K2P dimer. Prior to photolabeling experiments, purified TREK1 protein was assessed for functional integrity by reconstitution into planar lipid bilayers to measure single channel activity at the indicated holding potentials. Recordings were performed in symmetrical 150 mM KCl solution (D). hTRAAK protein was similarly active when reconstituted into bilayers (not shown).

**Supplemental Figure 2:**
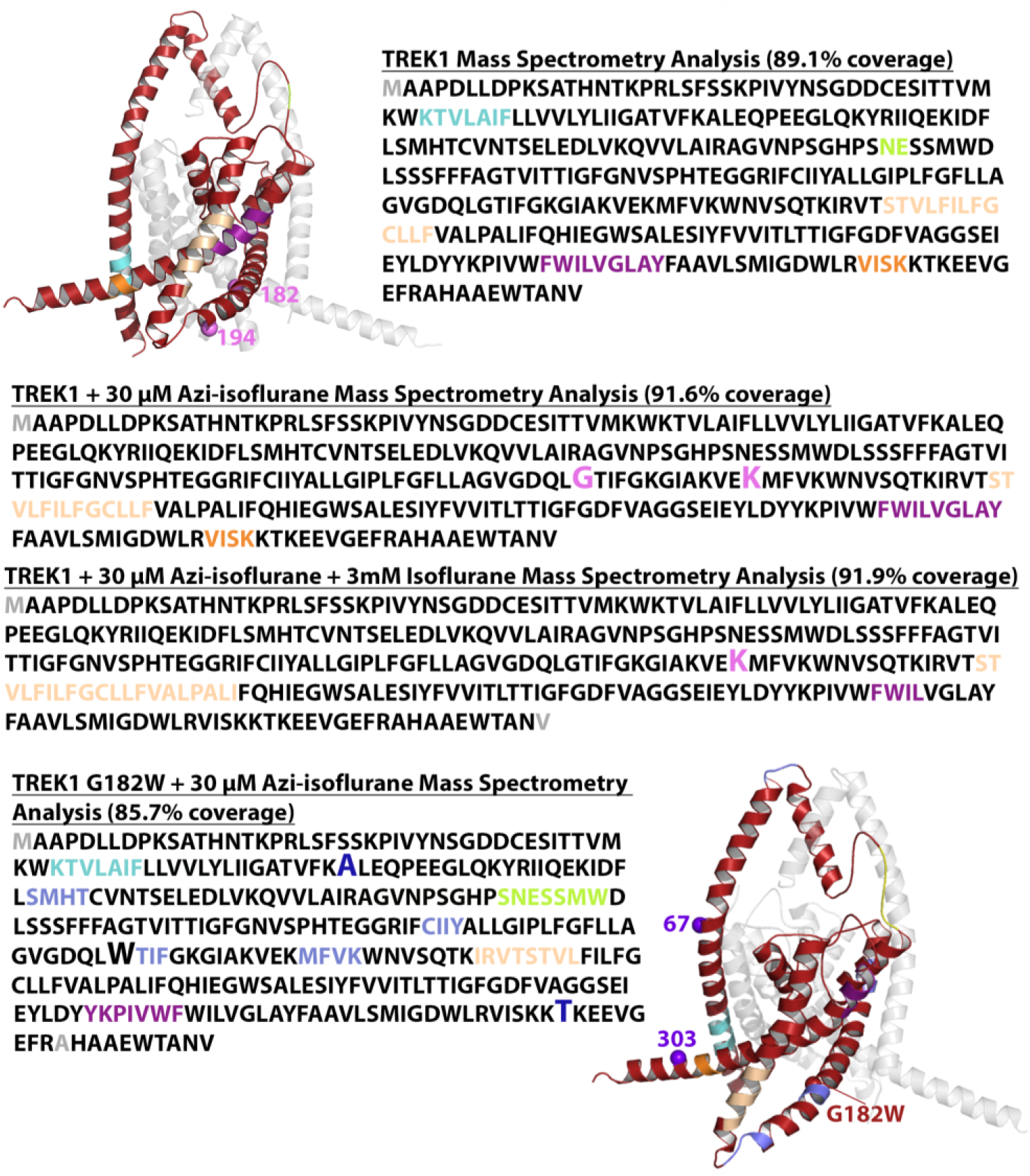
Mass spectrometry analysis of purified zebrafish TREK1. Results of MS analysis of TREK1 WT in the (top) absence of reaction with Azi-isoflurane, (second) following reaction with 30 μM Azi-isoflurane, (third) following reaction with 30 μM Azi-isoflurane in the presence of 3mM Isoflurane, or (bottom) TREK1 G182W following reaction with 30 μM Azi-isoflurane. Regions positively identified by mass spectrometry analysis are shown in red in the TREK1 structural model (PDB ID 6CQ6) and in black font in the sequence data. Regions absent from MS data occurred in five distinct regions, all of which are displayed in matching color in both the structural model and the sequence data. The G182 and K194 residues found to be modified by azi-isoflurane in TREK1 WT are shows as pink spheres in the structural model, and positive photolabeling is denoted in the sequence data by enlarged font and pink color. The A67 and T303 residues modified by azi-isoflurane in TREK1 G182W are similarly denoted in blue. The initial and final residues in the TREK1 protein were not identified in the majority of the MS results and are shown in grey to denote absence from positive MS identification. These residues are not present in the TREK1 structural model.

**Supplemental Figure 3:**
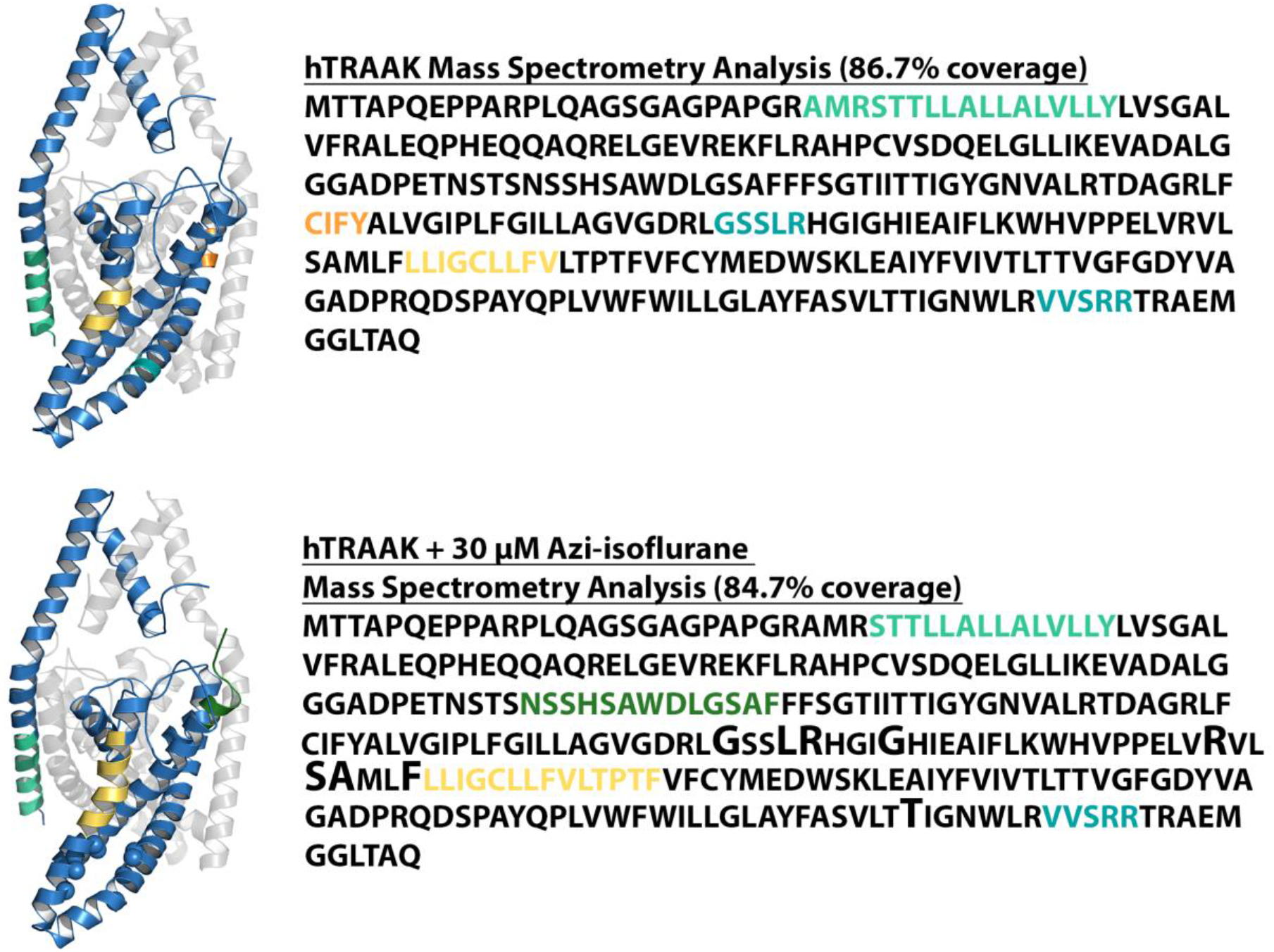
Mass spectrometry analysis of purified human TRAAK. Results of MS analysis of TRAAK in the absence of reaction with Azi-isoflurane (top) or following reaction with 30 μM Azi-isoflurane (bottom). Regions positively identified by mass spectrometry analysis are shown in blue in the TRAAK structural model (PDB ID 4WFE) and in black font in the sequence data. Regions absent from MS data are displayed in matching color in both the structural model and the sequence data. TRAAK residues homologous to TREK1 positions that exhibit high isoflurane occupancy in MD simulation are displayed as spheres in the structural model of TRAAK and are denoted in the sequence data by an enlarged font. All of these residues are identified by MS, but none show evidence of Azi-isoflurane labeling. A group of residues in the C-terminal region of TRAAK (marked in grey in the sequence data) were absent from our MS analysis but are not present in the TRAAK structural model.

**Supplemental Figure 4:**
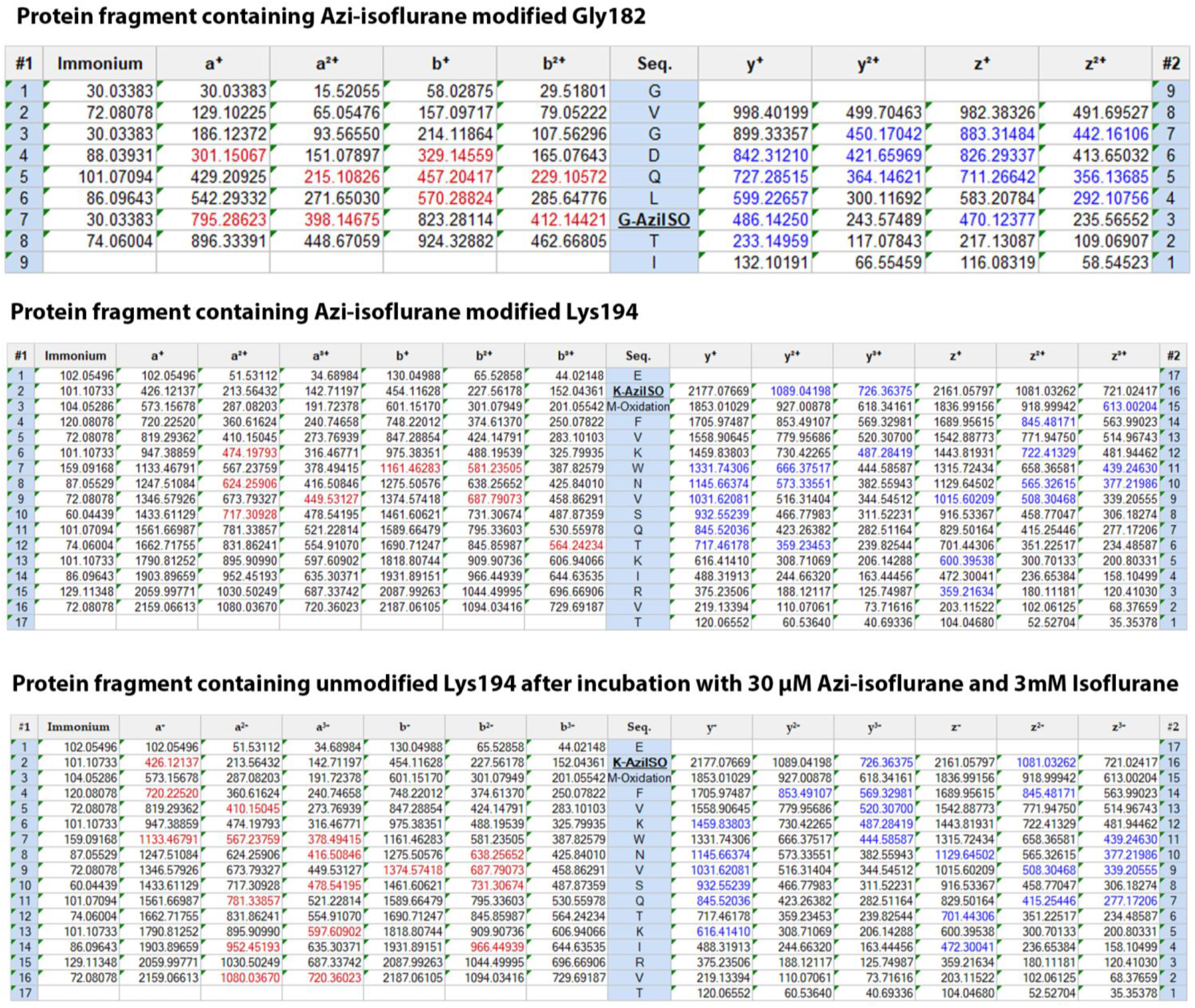
Mass spectrometry protein fragment tables. Shown are the fragmentation tables of the (top) 176-GVGDQL**G**TI-184 photolabeled peptide in the presence of 30 μM Aziisoflurane (middle) the 193-E<u>**K**</u>MFVKWNVSQTKIRVT-209 photolabeled peptide in the presence of 30 μM Aziisoflurane, and (bottom) the 193-E<u>**K**</u>MFVKWNVSQTKIRVT-209 photolabeled peptide in the presence of 30 μM Aziisoflurane (AziISO) and 3 mM isoflurane. Detected identified a, b (red) and z, y (blue) ions are colored red and blue, respectively. Residues detected with a modification are noted, and those modified by Aziisoflurane are additionally noted in bold and underlined.

